# PtrR (YneJ) is a novel *E. coli* transcription factor regulating the putrescine stress response and glutamate utilization

**DOI:** 10.1101/2020.04.27.065417

**Authors:** Irina A. Rodionova, Ye Gao, Anand Sastry, Jonathan Monk, Nicholas Wong, Richard Szubin, Hyungyu Lim, Zhongge Zhang, Milton H. Saier, Bernhard Palsson

## Abstract

Although polyamines, such as putrescine (Ptr), induce envelope stress for bacteria, they are important as nitrogen and carbon sources. Ptr utilization in *Escherichia coli* involves protein glutamylation, and glutamate stands at a crossroads between catabolism and anabolism. This communication reports that the transcription factor YneJ, here renamed PtrR, is involved in the regulation of a small regulatory RNA gene, *fnrS*, and an operon, *yneIHGF*, encoding succinate-semialdehyde dehydrogenase, Sad (YneI), glutaminase, GlsB (YneH), and several other genes. The *yneI* promoter is activated during putrescine utilization under nitrogen/carbon starvation conditions, and we show that PtrR is important for the putrescine stress response. It is also a repressor of *fnrS* gene expression, involved in the cascade regulation of mRNA synthesis for the *marA* and *sodB* genes, involved in antibiotic responses. PtrR transcriptional regulation of *fnrS* leads to a regulatory cascade induced by this small RNA that affects mRNA levels of *ompF* and the multidrug resistance regulator, MarA. We propose that PtrR functions as a dual activator/repressor, and that its regulation is important for the responses to different stress conditions involving L-glutamine/L-glutamate and putrescine utilization.

**IMPORTANCE:** Putrescine is an important source of nitrogen for many organisms, but it also induces stress. Although its metabolism has been studied extensively, the regulatory mechanisms that control the stress response are still poorly understood. This study reveals that the HTH-type transcriptional regulator, YneJ in *Escherichia coli*, here re-named PtrR, is important for the putrescine stress response, in part because it plays a role in outer membrane porin regulation as a sensor in a regulatory cascade. Direct PtrR transcriptional regulation of the *fnrS, yneI (sad), gltS* and *ptrR* genes is documented and rationalized, and nine PtrR binding sites were identified using ChIP-Exo. A *ptrR* mutant exhibited altered resistance to a tetracycline group of antibiotics under microaerophilic conditions, suggesting that PtrR indirectly controls expression of porin genes such as *ompF*.

## INTRODUCTION

*Escherichia coli* is a representative of the commensal mammalian intestinal microbiota and the best characterized model gram-negative bacterium. Nutrient starvation conditions are important for the gut microbiome bacterial community as they cause stress, activating different survival mechanisms (1). Many bacterial species, including *E. coli*, can simultaneously utilize L-glutamate (Glu) and the polyamine, putrescine (Ptr) under carbon/nitrogen starvation conditions, supporting bacterial growth using the PuuABCD pathway for Ptr utilization (Fig.1) (2). Glu is also essential for tetrahydrofolate polyglutamylation (3), for L-arginine biosynthesis, and for several other metabolic pathways. Ptr is important for bacterial growth and for efficient DNA replication, transcription, and translation (4,5). Organic polycationic molecules such as Ptr play important roles in maintaining compact conformations of negatively charged nucleic acids (6). However, high concentrations of extracellular Ptr are toxic for *E. coli*, altering the charge of outer membrane porins such as OmpF and OmpC (7,8), and Ptr is involved in multiple antibiotic resistance mechanism under stress conditions (7). The pore sizes decrease, resulting in pore closure, decreasing outer membrane permeability, and inducing stress response by activating Sigma E (Fig 1A). Recently, the transporter, AceI, and other members of the PACE family have been shown to function in the efflux of naturally occurring polyamines (9). These compounds have been described as key-signals for virulence in pathogenic bacteria, and they can support growth as nitrogen and carbon sources (10).

**Figure 1.**
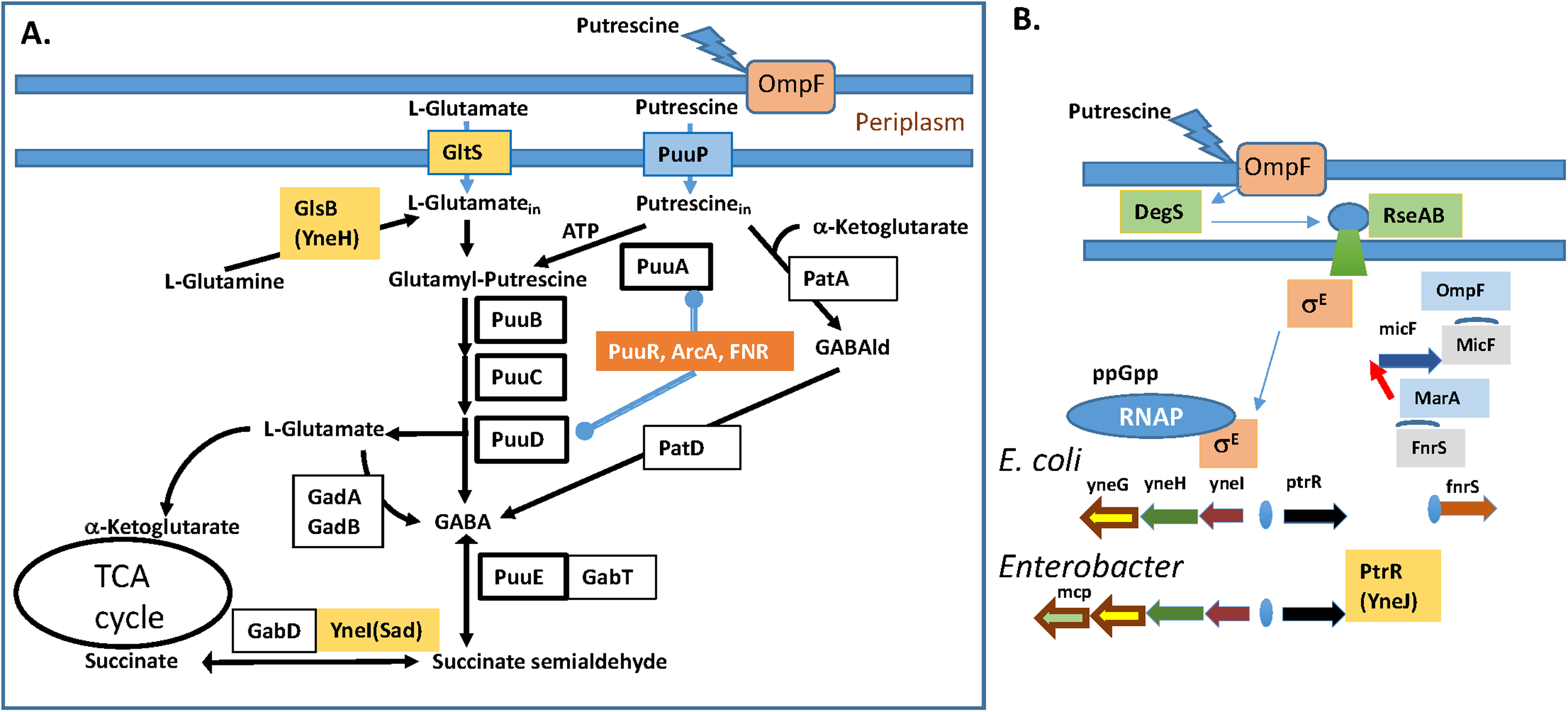
Regulation of putrescine utilization and the response to induced stress. **A.** The putrescine utilization pathways. PtrR (YneJ) regulated genes are shown in yellow boxes. Abbreviations: GABAld – gamma-aminobutyraldehyde, PatA - putrescine aminotransferase, PatD - gamma-aminobutyraldehyde dehydrogenase, YneI (Sad) - succinate-semialdehyde dehydrogenase, YneH (GlsB) - glutaminase. The *puu* operon is regulated by the PuuR, ArcA and FNR transcription factors. **B.** The PtrR (YneJ) regulation of *fnrS* and the cascade regulation for *ompF* as well as Ptr utilization. The cascade regulation of MarA mRNA by fnrS and MarA transcriptional regulation of micF - small RNA (sRNA), regulating OmpF mRNA stability.

In bacteria, the processes of Ptr uptake, synthesis, degradation, and export are coordinated to stringently regulate intracellular polyamine levels. Here, we demonstrate by ChIP-exo assays that the LysR-family transcriptional regulator, YneJ (PtrR), binds to the promoter regions of at least eleven genes. A *yneJ* mutant was shown previously to be resistant to bacteriophage lambda infection, probably because it transcriptionally regulates *lamB* (maltoporin)(11). RNA sequencing analyses suggested that PtrR under carbon starvation conditions with the addition of L-glutamine and Ptr repressed the *sad* (*yneI)* and *fnrS* genes (Fig. 1A), encoding succinate-semialdehyde dehydrogenase (SSADH) and a small RNA. In addition, the *yneI* operon encodes glutaminase (YneH), and two hypothetical proteins, YneG and YneF. The regulatory small RNA, FnrS, is known to regulate mRNA stability for the transcriptional activator of the multiple antibiotic resistance gene *marA* (12). FnrS is regulated by Fnr and ArcA (13). The transcriptional regulation of FnrS by PtrR is likely important for the Ptr-induced stress response.

Ptr utilization is catalyzed by enzymes encoded by the *puuABCDE* operon in *E. coli* and involves degradation of putrescine to γ-aminobutyric acid (GABA) via γ-glutamylated intermediates (Fig. 1A). PuuA, glutamate-Ptr ligase, the first enzyme in the pathway, and is regulated at the transcriptional level by PuuR, Fnr, and phosphorylated ArcA (2). Although the putrescine pathway enzymes PatA (YgiG) and PatD (YdcW) comprise a pathway for the degradation of putrescine without glutamylation (14), the PuuABCDE pathway is essential for Ptr utilization in *E. coli* using PuuP as the major transporter (Fig. 1A) (15).

It is not entirely clear why *E. coli* and many other bacteria have two genes encoding 4-aminobutyrate aminotransferases (GABA-ATs) (GabT and PuuE or GoaG) as well as two succinate semialdehyde dehydrogenases (SSADHs) (GabD and YneI or Sad)(16,17). The explanation may have to do in part with the fact that regulation of *gabDT* and the GABA transporter gene, *gabP*, serves for GABA utilization. The *csiD-ygaF-gabDTP* region is sigma S (σ^S^) controlled (18), and activation of an internal promoter within *gabD* is also under the control of σ^S^ (19). GabT function has been well studied (19–21).

Expression analyses of PuuE and YneI have also been reported (22,23), but as noted above a transcriptional regulator for the *yneI* gene had not been found. PuuE is important for GABA metabolism in *E. coli* and is induced by the addition of putrescine to the medium. It is repressed by succinate under low aeration conditions, although in contrast, the *gabT* expression level is not influenced under these conditions. However, under respiratory stress conditions, other data suggest dramatic up-regulation of GabT and GabD as well as genes for respiration, the glyoxylate shunt, and motility (24). Moreover, YneI is induced by putrescine in the medium while GabD is not. In *E. coli*, both PuuE and YneI are important for the utilization of putrescine as a sole carbon source. GABA utilization-related YneI (the structure of which is known (25)) has been shown to be essential for the utilization of putrescine as a carbon source (22). However as noted above, GABA utilization enzymes are redundant in *E. coli*; the genes for SSADH are *gabD* and *yneI*, while those for the GABA aminotransferases (GABA-AT) are *gabT* and *puuE*. The GABA shunt may have pleiotropic functions under different stress conditions (21).

In this study, we investigated PtrR-dependent regulation, important for the Ptr stress response and for Ptr utilization by the coordination of *yneIHGF, fnrS*, and likely *gltS* transcription. The binding sites upstream of *yneI*, *ptrR* (*yneJ*), *gltS*, and *fnrS* were identified using the ChIP-exo method, and were used to determine the consensus predicted, based on the alignment of the *yneI* and *fnrS* or *ptrR* and *fnrS* upstream sequences for the PtrR ChIP-exo protected areas (Fig. 2A and B). In fact, a total of nine binding sites with different intensities for PtrR were identified (Table 1, Fig. 2B), and our experiments revealed regulation by the binding of PtrR upstream of the *ptrR, yneI* and *fnrS* genes. It is likely, that PtrR transcriptionally regulates the *ydiFO* genes, which encode acetyl CoA transferase and acyl-CoA dehydrogenase involved in the fatty acid degradation pathway (26). PtrR is therefore a novel pleiotropic LysR family transcriptional factor, and it represses transcription of *fnrS* and *yneIH*. The activity of the *yneI* promoter has been shown to increase upon addition of Ptr to the growth medium under nitrogen starvation conditions (Fig.2C), and a *ptrR* mutant showed a growth defect in M9 medium supplemented with Glu as a sole nitrogen source and glycerol as a carbon source. The positive effect for the growth of *ptrR* and *ydiF* mutants on Ptr and Glu as nitrogen sources under carbon starvation was also shown. We conclude that PtrR controls the Ptr stress response and Glu utilization in *E. coli*, probably in part in response to GABA availability. The degraded palindromic consensus for the PtrR binding motif for the *fnrS* and *yneI(sad)-yneJ* genes was determined. The predicted PtrR regulons in other bacteria revealed novel MFS transporters possibly related to GABA utilization.

**Table 1.**
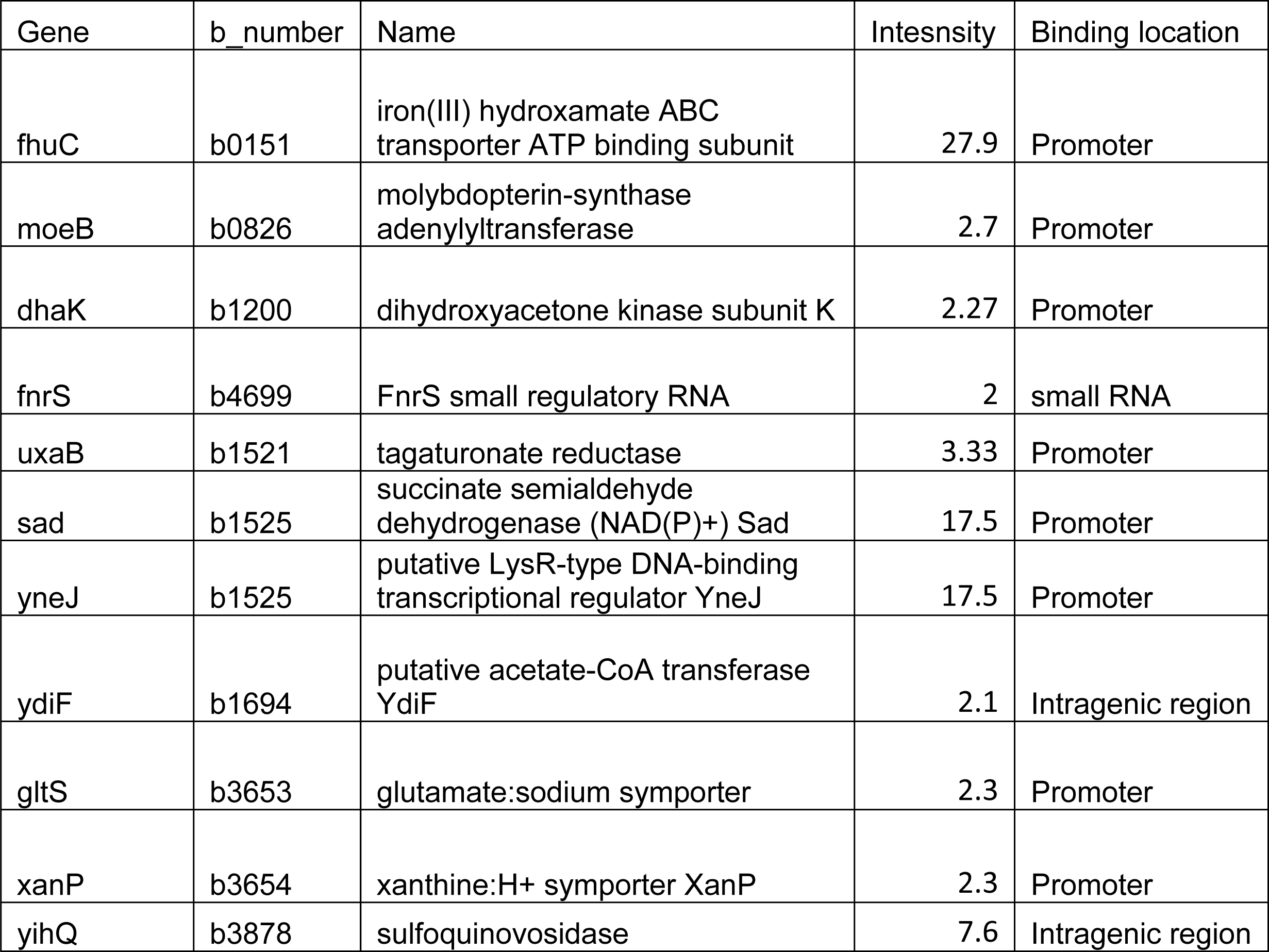
ChIP-exo detected PtrR binding sites in the promoter regions and intergenic regions of *E. coli* genes.

**Figure 2.**
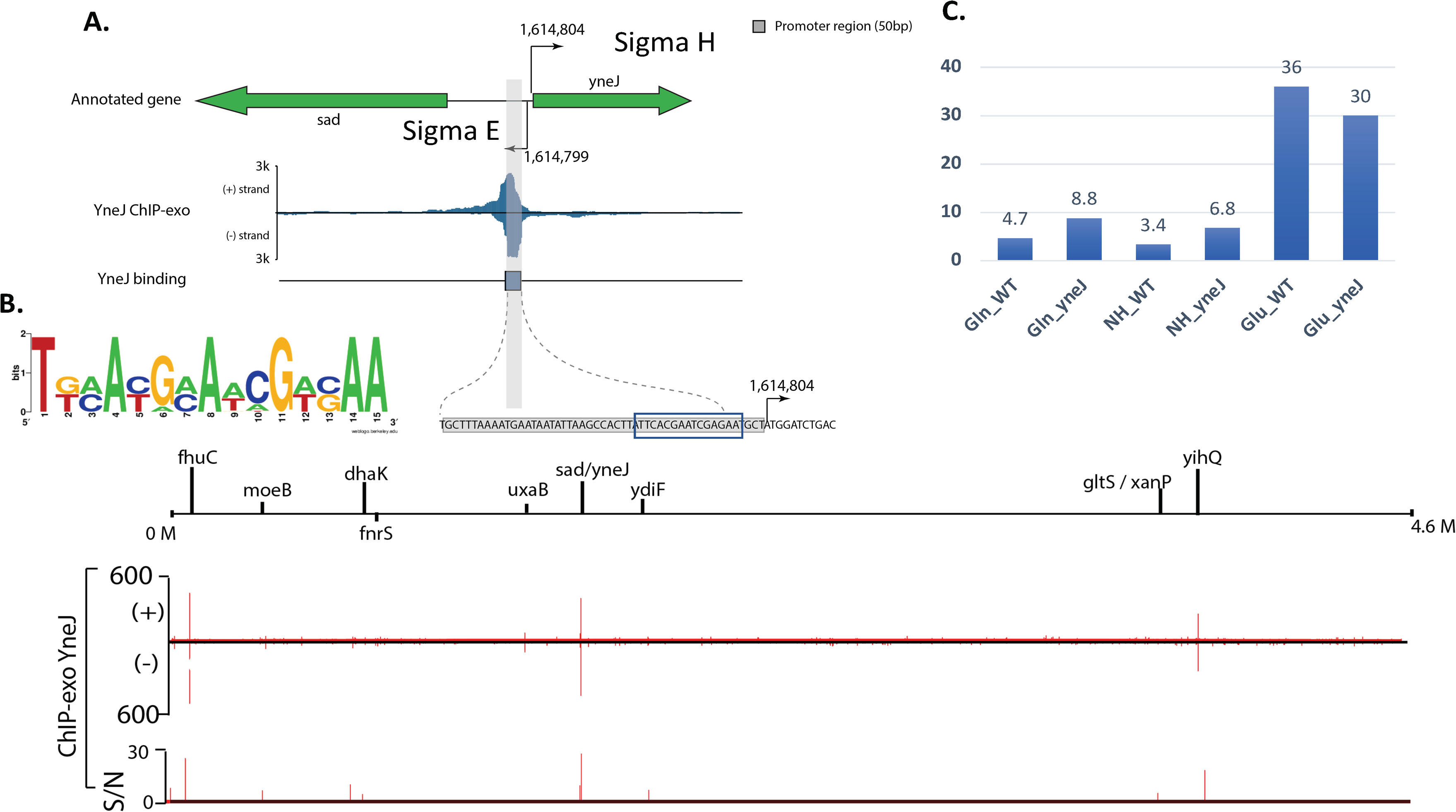
*yne* promoter regulation. **A.** The *yneJ* and *yneI* promoter region for PtrR (YneJ) binding, identified by ChIP-exo, **B.** The predicted consensus in the WT BW25113 strain sequence, and the corresponding consensus with the *yne* operon from other proteobacteria. **C.** The promoter activity measured by -galactosidase assay for the WT-wild type and *yneJ* mutant strains grown for 3 hours in M9 medium with 0.4% glycerol and the nitrogen source as marked: either NH-NH_4,_ or Gln-glutamine, or Glu-glutamate with 10 mM Ptr added.

## RESULTS

### Predicted regulatory PtrR binding sites and ChIP-exo results

The putative binding site for PtrR, located close to the Sigma E (σ^E^)-dependent promoter region of *yneI*, was predicted using a bioinformatics approach and confirmed by ChIP-exo results (Fig.2A and B, Table 1). The sequence logo for identified PtrR-binding sites from the intergenic region of the *yneJ* (*ptrR*) and *yneI* genes from *E. coli* and closely relates enterobacteria, as well as similar DNA sites identified upstream of the *fnrS* genes is shown in Fig. 2B. The *yneI-sad* operon is well conserved in Enterobacteria, but in *Enterobacter*, the operon includes an additional gene for the methyl-accepting chemotaxis protein I (serine chemoreceptor protein, Mcp) (Fig. 1B). A bioinformatics approach for consideration of the co-regulation of the *ptrR* and *puuABCDE* genes revealed that *ptrR* is present in the genomes of organisms that lack the *puuABCDE* genes. In these organisms, the pathway for Ptr utilization without glutamylation involves PatAB (Fig.1A), suggesting that regulation of the polyamine stress response via *fnrS* gene repression is an additional function of the PtrR regulator.

PtrR regulatory binding sites were identified by ChIP-exo. PtrR binds upstream of the *fnrS*, *gltS*, *ptrR* (*yneJ*), and *yneI* genes, and in an intragenic region of *yhiQ* and *ydiF*. It is also likely to be involved in its own autoregulation. PtrR (YneJ) binding sites for the *gltS* gene were found to be upstream of the promoter (Fig. S1). The *yne* operon appears to be regulated by the stress sigma factor E in addition to PtrR.

### *E. coli* growth on glutamate as a nitrogen source and an additional carbon source

Growth of BW25113 (WT) as well as *ptrR* null mutant strains could occur in M9 medium with glucose or glycerol as the carbon source and Glu as the nitrogen source, but not with Glu as the sole carbon and nitrogen source. Cells showed a growth deficiency when the *ptrR* gene was deleted under nitrogen and carbon starvation conditions with both Glu and glycerol present in minimal medium (Fig. 3). A decrease in the growth rate was observed on glycerol (carbon starvation) for the *ptrR* mutant (Fig 3), suggesting that activation of the *gltS* promoter is essential for rapid growth. Substantiating this possibility, the *gltS* null mutant was growth deficient in M9 medium to the same extent under these defined conditions. However an mRNA level change for *gltS* in the *yneJ* mutant was not detected; instead, the *yneI* mRNA level was higher in the *ptrR* mutant. This result suggests that the flux through YneI (Sad) was increased in the *ptrR* mutant, probably decreasing Glu input as a nitrogen/carbon source. Strong upregulation of the *sad* gene in the *ptrR* mutant relative to the WT *E. coli* BW25113 strain was observed by mRNA level measurements during growth on Glu as the single nitrogen source and glycerol as the carbon source (Table S1). The *ptrR* mutation led to the upregulation of L-histidine and L-methionine biosynthetic pathway genes *hisABCDFHI* and *metABEFLR*, respectively. Additionally, upregulation of more than a hundred genes including *fliA, ddg, dppBCDF, cspA*, *ybdHL*, and *ykfD* observed in the *ptrR* mutant.

**Figure 3.**
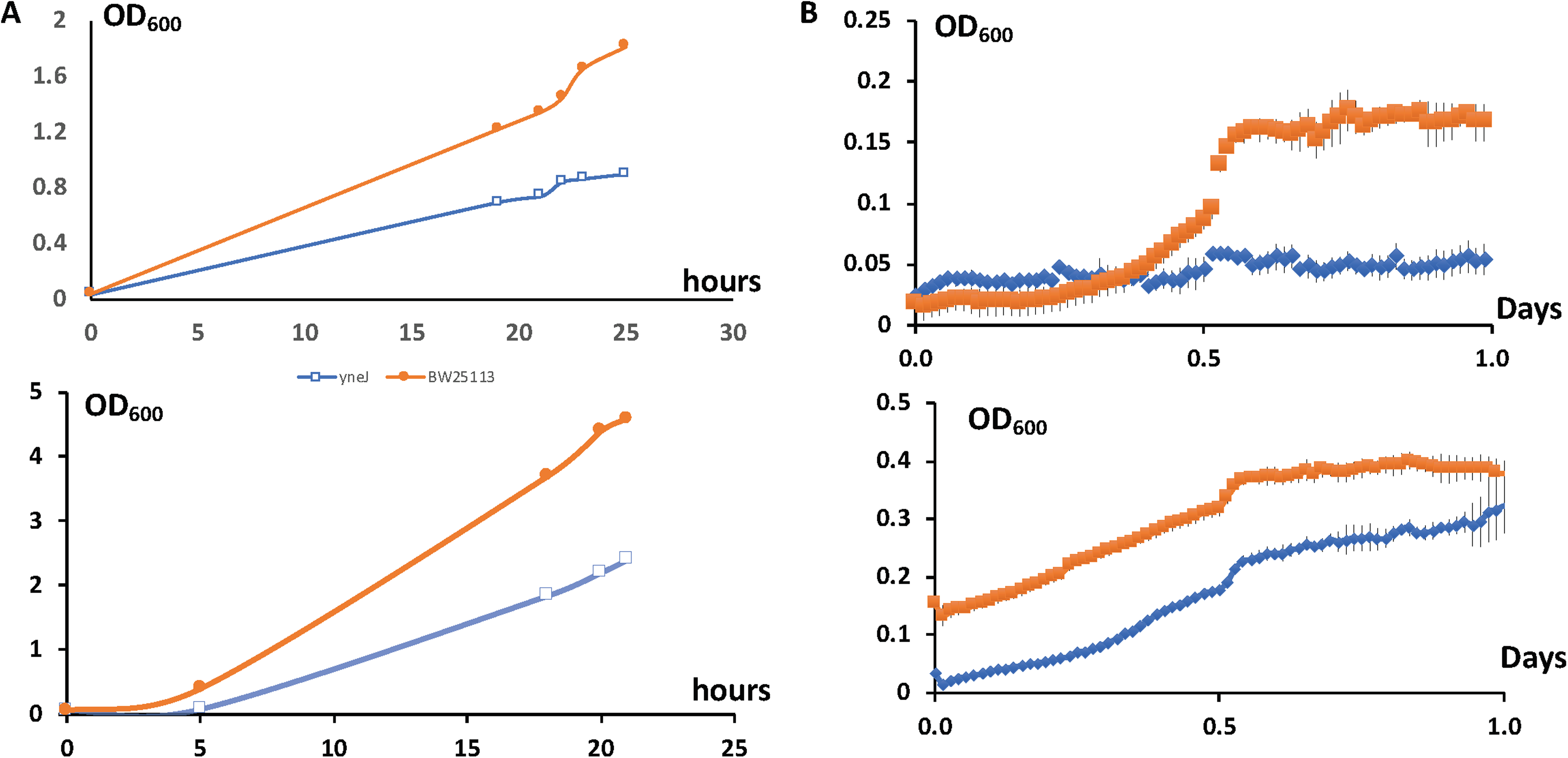
Growth of the *ptrR (yneJ)* mutant (blue marker) compared to the WT BW25113 strain (red marker) with 20 mM L-Glu as the sole nitrogen source and glycerol as the carbon source, **A.** Growth in overnight tubes. The glycerol concentration was 0.2% (upper) or 0.8% (lower). **B.** The WT BW25113 and *ptrR* mutant strains were grown under the same conditions with 0.2 % glycerol in 96-well plates (upper) or in normal glucose-M9 medium (lower).

CspA is a cold shock protein regulating many genes, likely related to starvation conditions. The RNAseq analysis using the recently developed independent component analysis (ICA) showed that the modulon (27) for *cspA* includes *lpxP*, the transferase involved in KDO-lipid A modification under cold shock conditions. We showed that under carbon starvation, especially for the *ptrR* mutant, the CspA modulon genes were highly induced. The enterobacterial common antigen biosynthetic genes, *wecCD* and *rffH*, were also upregulated in the mutant.

Growth suppression for the *ptrR* mutant was observed under microaerobic conditions in 96-well plates with L-glutamate as the nitrogen source and glycerol as the carbon source (Fig. 3). In contrast, no difference was detected in the growth rate for the *ptrR* mutant when cells were grown on glucose as the carbon source with glutamate as the nitrogen source (data not shown). We tested the microaerobic conditions using Biolog plate PM2A for *ptrR* mutant growth compared with that of the *E. coli* WT BW25113 strain. The difference in growth rate using Glu as the nitrogen source was minimal when D-glucosamine, dihydroxyacetone or D-lactate methyl ester was the carbon source. Poor growth of the mutant with glycine, 5-keto-gluconate, L-ornithine, or gamma-hydroxybutyrate as a carbon/nitrogen source was observed using M9 medium with L-glutamate as the nitrogen source.

Deficiencies of *ptrR* mutant growth with D-allose, D-tagatose, turanose, oxalomatic acid, methyl-glucuronide, gamma-hydroxybutyrate, glycine and L-alaninamide were detected under the same conditions with Glu as a supplement (Fig S2, Table S2). A small difference for growth with D-arabinose and 5-keto-gluconate was detected using the same conditions. No difference in growth rate was observed when dihyroxyacetone, D-lactate methyl ester, or melibionate was the carbon source.

### The regulatory effect of PtrR during growth with putrescine or Glutamate as the nitrogen source

M9 medium with glycerol as the primary carbon source, along with the Ptr and Glu both present as carbon/nitrogen sources, supported the growth of BW25113 (WT) as well as *ptrR, yneH*, and *ydiF* null mutant strains (Fig. 4). The WT strain had a longer lag-phase compared to the *yneJ* mutant, possibly because transcriptional regulation plays a role in normal growth. The presence of Ptr as a nitrogen source induced the *yne* operon during nitrogen/carbon starvation. Derepression of the operon occurred in the *yneJ* mutant, which led to faster adaptation to Ptr. The importance of *yne* operon expression under M9 carbon/nitrogen starvation conditions with Ptr as a nitrogen source was demonstrated. A growth deficiency for a *yneH* mutant was observed under the same conditions. The effect of a *yneH* deletion was strongest as cells approached the stationary phase. This result is interesting because Sigma E-dependent promoters have been shown to be induced in stationary phase (28).

**Figure 4.**
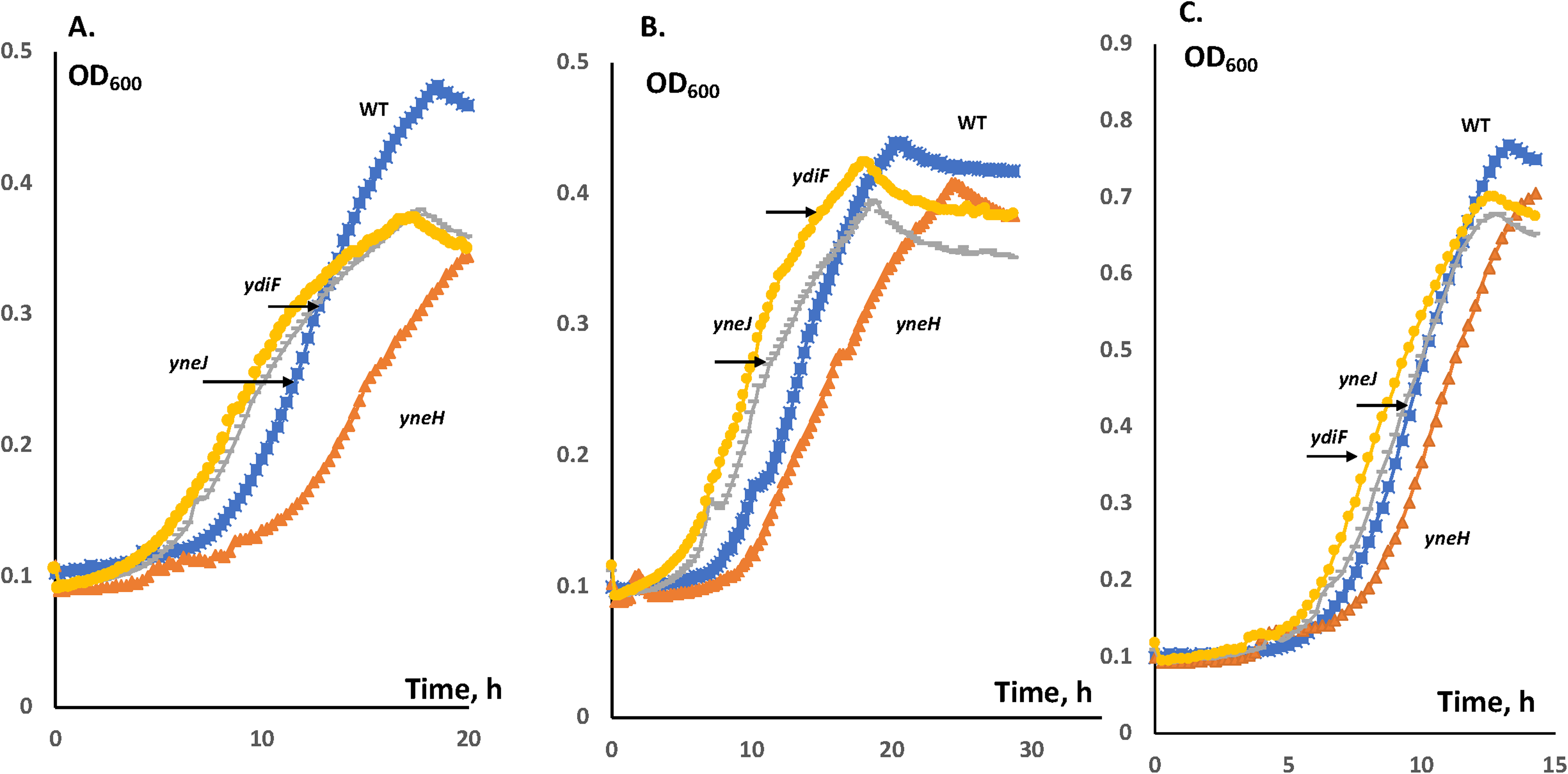
Growth of *yneJ*, *yneH* and *ydiF* mutants compared to the WT BW25113 strain **A.** M9 medium with 20 mM Glu and 10 or 20 mM Ptr, as nitrogen sources and 0.4% glycerol (v/v) as the primary carbon source. **B.** The cells were grown under the same conditions, but 20 mM Ptr was added. **C.** The cells were grown in M9-glucose medium.

The *ydiF* mutant phenotype is shown in Fig.4. A PtrR binding site was found inside the *ydiF* gene. *ydiO*, downstream of *ydiF*, but within the operon with *ydiF*, encoding acyl-CoA dehydrogenase, is potentially affected by PtrR at low concentrations of Ptr, and an *ydiF* null mutant had a similar phenotype as that of the *yneJ* mutant. It is possible that YdiF-YdiO interferes with the usage of succinate, produced from Ptr. This could reduce the succinate pool, which is important for fatty acid biosynthetic fluxes, negatively affecting metabolic pathways during growth on Ptr as an additional carbon source. The *ydiO* gene has been shown to be important for the utilization of fatty acids in a *fadE* mutant of *E. coli* (26).

### PtrR-dependent regulation during growth with Ptr/Glu as nitrogen sources

The *E. coli* WT and the *yneJ* mutant were grown in M9 medium with 20 mM Glu and 8 mM Ptr as nitrogen sources and 0.2 % glycerol as the primary carbon source. The growth phenotypes are shown in Fig. 3. To determine the effect of the *ptrR* deletion mutation, the cells were collected at the end of log-phase, and total mRNA was purified (see Materials and methods). PuuR and PuuADE mRNA levels increased in the *ptrR* mutant strain, possibly explaining the faster growth of the mutant under the experimental conditions, but lower final OD_600_ in the stationary phase (Fig.2A). Although the binding of PtrR in the *puu* operon promoter area was not demonstrated using the ChIP-exo method, the effect of deletion of *yneJ* on Ptr utilization gene expression (direct or indirect through derepression of *yneI*) defined PtrR as a Ptr related transcriptional regulator. It is interesting that two copper related transport systems were upregulated in the *ptrR* mutant. These genes encode CopA (copper efflux system) and the CusSR regulated CusABC system (a cation efflux system), shown as copper I-modulon, additionally including *cusF and cueO*. The FnrS and RyhB small RNAs influence a regulon that includes SodB (superoxide dismutase), as evidenced by their binding to the SodB mRNA (see EcoCyc). SodB, RpmL, YihD, and PuuRADE mRNAs were substantially higher due to the loss of *ptrR* compared to the WT strain (Table 2). It is likely that the effect of *fnrS* transcriptional derepression, due to the *ptrR* mutation, led to an increase in the SodB mRNA level under these experimental conditions.

**Table 2.**
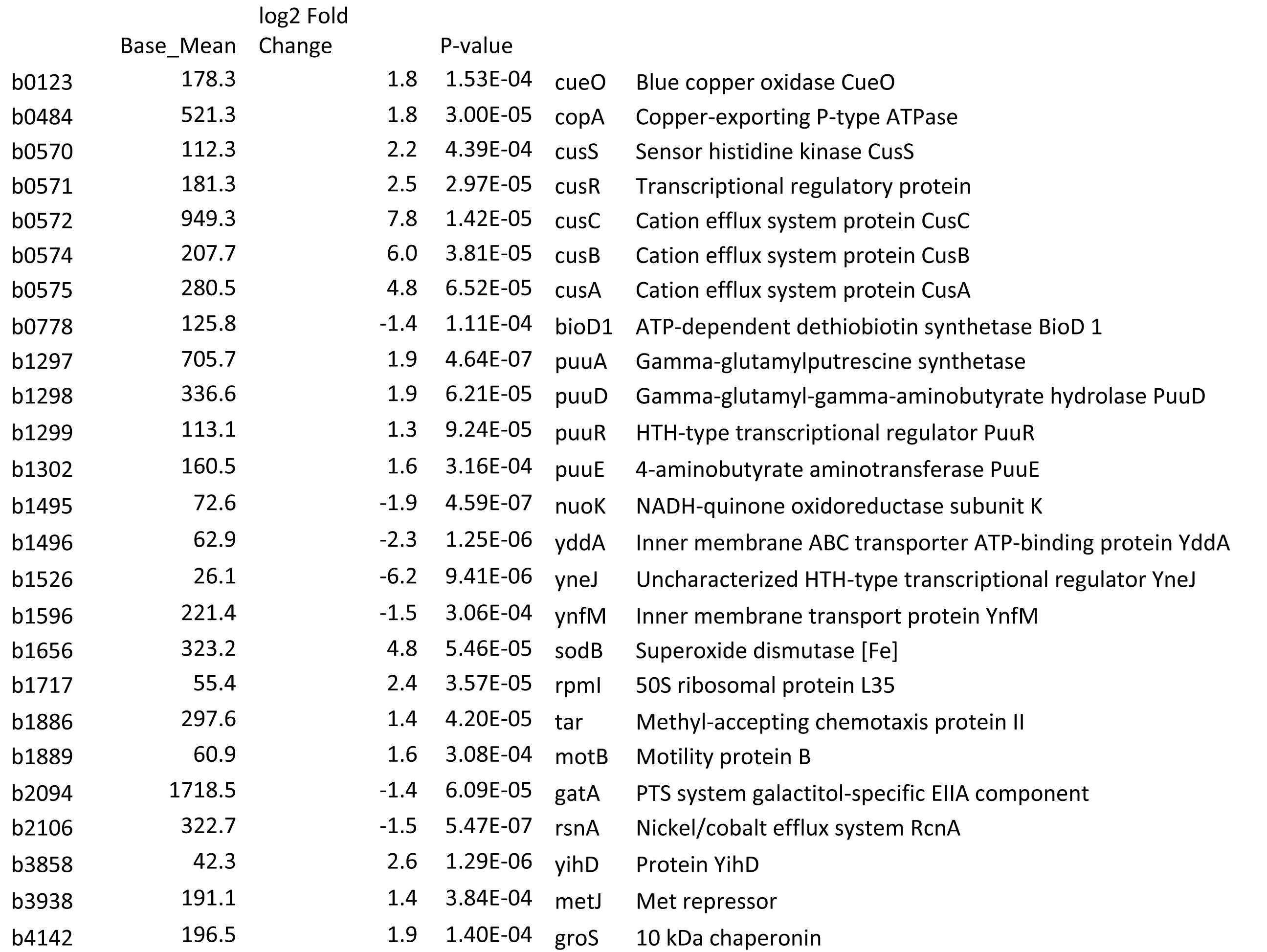
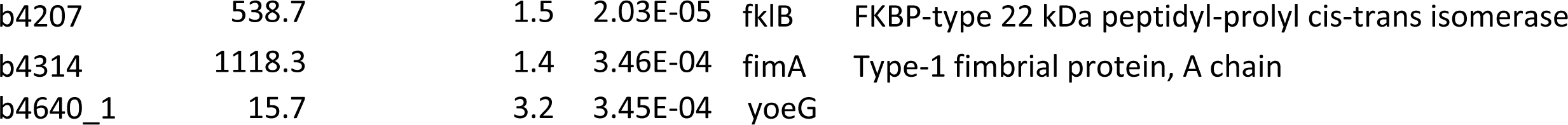
Differentially expressed genes revealed by the RNAseq data for a *ptrR* mutant and WT *E. coli* BW25113 strain during growth in M9 medium with L-Glutamate and Putrescine as nitrogen sources and glycerol as the primary carbon source

### PtrR-dependent regulation of the *fnrS* and *ompL* genes, and *yne* operon expression under putrescine stress conditions

The *ptrR* mutant and the wild type BW25113 strain were grown aerobically in M9 medium under carbon starvation conditions (glycerol as the carbon source), supplemented with 20 mM L-glutamine and 10 mM Ptr. RNA sequence data were generated during the exponential growth phase. The effects of putrescine on the *ptrR* mutant were determined by RNA-seq analysis. The mRNAs whose levels were depressed in the WT strain compared to the *ptrR* mutant are *yneH* (*glsB*), *yneI* (*sad*), and *yneG*. The mRNA levels for *acrS, tolC, yihO, ompL* and *ompF* were higher in the WT strain (Table S3), suggesting a regulatory cascade for PtrR-FnrS-MarA-MicF-OmpF and PtrR transcriptional activation of the *yihPO-ompL* operon. MarA alters the expression of more than sixty chromosomal genes (29), and efflux/influx-related genes are important for the MarA-related phenotype. ChIP-exo showed PtrR binding in *yihQ* gene, but only part of operon (*yihO-ompL*) mRNA was found to be differentially expressed. It is likely that PtrR is activator for the *yihPO-ompL* and/or mRNA degradation is involved in the mRNA homeostasis. The operon is conserved in Enterobacteria as shown by use of the PubSeed database (Fig. S3).

We decided to test the *ptrR* mutant for antibiotic resistance using Biolog plate 11C (30). This regulatory mutant showed increased resistance to high concentrations of demeclocycline, a tetracycline group antibiotic, which showed growth/respiration after 3 days, but poor growth/respiration compared to the *E. coli* WT strain in the presence of certain concentrations of lomefloxacin, cloxacillin, colistin, and potassium tellurite, but no difference was observed with respect to the resistance to other antibiotics tested. The resistance of the *ptrR* mutant, also to the tetracycline analog, chlorotetracycline, suggests that *ompF* mRNA was downregulated under these experimental conditions. The role of the outer membrane barrier for efflux-mediated resistance has been discussed (31). MarA activates *tolC*, *zwf, micF, ybjC, sodA, inaA*, and *mdaB*, but the small RNA, MicF, specifically inhibits *ompF* mRNA translation (32).

### Promoter activity for *yneI* under nitrogen starvation and Ptr-induced stress conditions

The activity of the promoter for the *yneI* gene was measured during growth in M9 medium with Glu or L-Glutamine as the sole nitrogen source and glycerol as the carbon source, but no activation was observed (data not shown). In contrast, under the same conditions, but with Ptr present, the promoter activity for *yneI* increased steadily under nitrogen starvation conditions with Glu as nitrogen source (Fig.2B). The activation effect was not detected in glycerol-M9 medium with NH_4_Cl as the nitrogen source.

We found that PtrR regulates the outer membrane porin encoded by the *ompF* gene (likely via FnrS) as well as the *yne* operon (Table S3). Transcription of the *E. coli yneIHGF* operon was induced when Ptr was present under nitrogen starvation conditions. The operon encodes Glu/Ptr catabolic pathway enzymes and is conserved among many Gram-negative bacteria (Fig. 1). The PtrR regulon includes the *yneIHGF* operon and the *gltS* gene as a part of the putrescine utilization pathway since PtrR regulates the promoter of the *yneI* gene under Ptr-induced stress conditions. As noted above, the *yne* operon includes *yneI* (*sad*) and *yneH* (*glsB*), and repression of *sad* was shown by RNA-seq data. The Glu transporter gene, *gltS*, is likely activated by PtrR, which could be important for *E. coli* growth with Glu as a nitrogen source and glycerol as a carbon source (nutrient starvation conditions). YneI (Sad) is known to be involved in Ptr utilization (22). Using the ChIP-exo method, we showed that PtrR has nine binding sites in *E. coli* for genes including the glutamate transporter gene, *gltS*, the regulatory sRNA gene, *fnrS* and *yihQO*. Phenotypic analysis and RNAseq results for a *ptrR* deletion mutant confirmed that transcriptional regulation of the *yneIHGF, fnrS* and possibly the *yihOP-ompL* was important during Ptr and glutamine utilization and stress response.

### Prediction of PtrR binding consensus for *fnrS* and the *yneI*-*yneJ* genes

We created multiple alignments of the upstream DNA sequences of closely related species with the beginning of the *E. coli* gene for *fnrS*, as well as upstream of the *yneI* and *yneJ (ptrR)* genes. These binding sites correspond to the ChIP-exo protected areas. The binding sites TTCACGAATCGaGAA, TTCtCGATTCGTGAA, and TgaAtGcAaCGTcAA were found for *yneJ (ptrR)*, *yneI*, and fnrS, respectively (Fig. 5). The fluorescent polarization assay showed binding for PtrR refolded protein in the presence of 10 mM urea to the predicted *yneI* binding site. The addition of GABA lead to dissociation of the PtrR from fluorescently labeled DNA (Fig. S5). The PtrR potential binding sites for *yihO* and *yaiYZ* encoding transporters and *asnB*, asparagine synthetase B and uncharacterized protein, *ydcY*, were predicted and confirmed by RNAseq data (Tables S1, S3). ydcY gene is conserved in genome cluster with *potABCD* encoding ABC type transporter and gamma-aminobutyraldehyde dehydrogenase gene, *patD*, involved Ptr utilization (Fig.1). The PtrR binding sites for species of *Pseudomonas, Salmonella, Yersinia, Serratia, Klebsiella, Enterobacter, Citrobacter, Erwinia*, and *Shigella* were also identified, predicting *yne* operon regulation similar to that observed for *E.coli*. The predicted regulatory sites for MFS family transporter, likely related to GABA or succinate semialdehyde uptake in seven *Yersinia*, three *Serratia*, one *Erwinia carotovora*, and one *Pseudomonas aeruginosa* genomes were identified. In some of these genomes, the regulons are the same as the *E. coli yne* operon and include *yneI* and *yneH*, but binding site for *fnrS* was not found. In *Pseudomonas*, and many of these γ-Proteobacteria the *yneI*(*sad)* gene is regulated by PtrR, but in *Enterobacter sp. 638* and *Citrobacter koseri* additional genes are included (Fig. 5).

**Figure 5.**
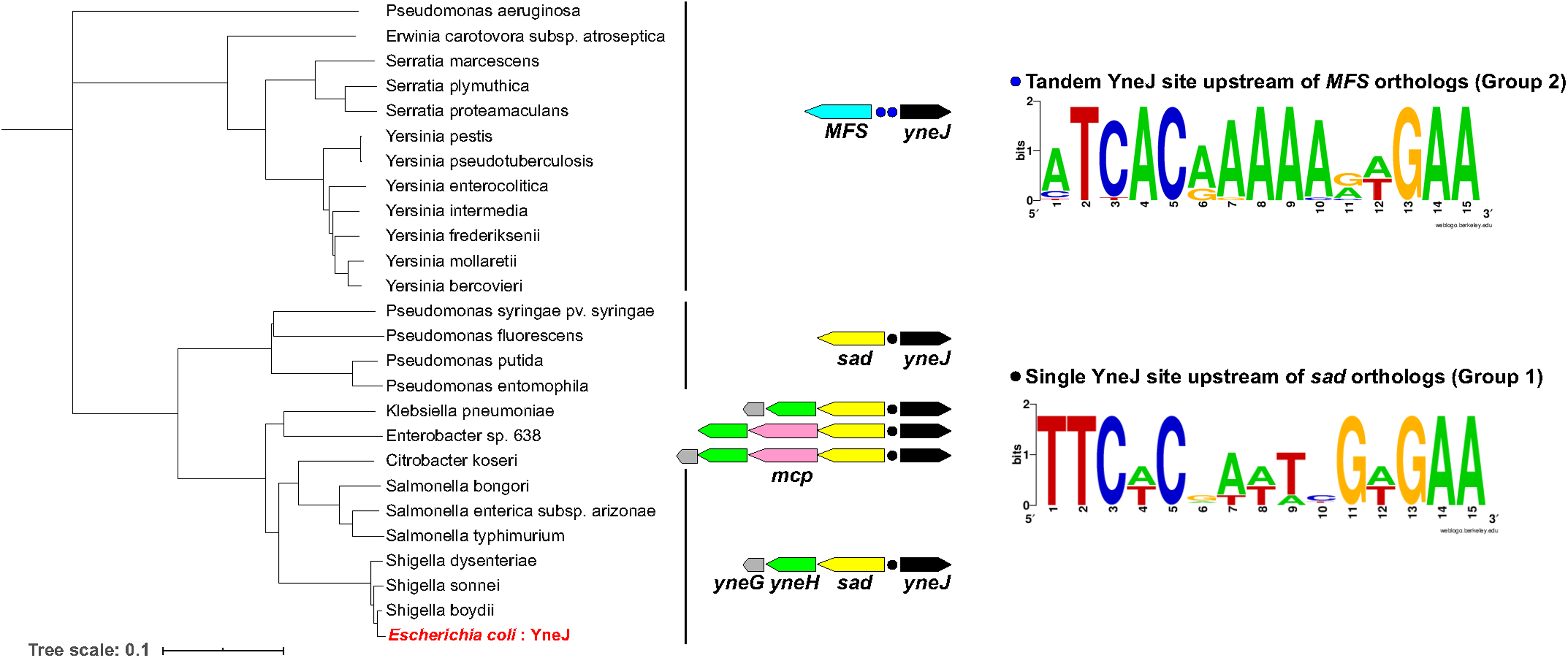
The PtrR phylogenetic tree and the predicted binding consensus sequence for PtrR in other Proteobacteria. The regulons for several Proteobacterial representatives are shown.

## DISCUSSION

In this study, we have characterized PtrR (YneJ) as one of the transcriptional regulators for operon encoding glutaminase, the GABA shunt SSADH, YneI (Sad), GlsB (YneH), and the uncharacterized proteins, YneG and YneF, as well as *yihPO-ompL* operon, the gene encoding the glutamate transporter, *gltS*, and the small RNS, FnrS. We found that PtrR is important for the Ptr and L-glutamine stress response and utilization of L-glutamate as a sole nitrogen source with glycerol as the carbon source under the carbon starvation stress conditions used in our studies. We further demonstrated PtrR binding to the predicted DNA-binding site using the ChIP-exo method. An increase in the promoter activity for *yneI* under putrescine stress conditions with either nitrogen or carbon starvation was found to be PtrR dependent. PtrR is a repressor for the *yneIHFG* operon under these stress conditions.

Based on the growth phenotypes, PtrR-mediated regulation appears to be important for Glu utilization as a nitrogen source in the presence of a poor carbon source such as glycerol. A pleiotropic effect of the PtrR-dependent regulation of the sigma E-dependent *yneIHGF* operon under nitrogen/carbon starvation is related to Ptr-induced envelope stress conditions. Analogous to the known stress/starvation sigma σ^S^-controlled *csiD-ygaF*-*gabDTP* region (15,19), the *yne* operon is dependent on the environmental sigma factor σ^E^. Growth of the *ptrR* mutant with Glu as the sole nitrogen source and glycerol as the carbon source allows production of ppGpp (33,34).

Ptr is involved in both envelope stress and starvation stress, and the importance of PtrR regulation of the *yne* operon under these stress conditions was shown by assay of β-galactosidase. The *yne* promoter was highly upregulated under carbon/nitrogen starvation conditions with 20 mM Ptr and 20 mM Glu as nitrogen sources or with Glu as a nitrogen source (RNA-seq data) and glycerol as a carbon source (Fig. 2). The growth deficiency of the *ptrR* mutant suggested that PtrR-dependent activation of the *gltS* promoter occurs under nitrogen starvation conditions. However, a difference in mRNA level was not detected. PtrR autorepressor function was suggested from the binding site position, downstream of the *ptrR* promoter (Fig. 2). The *ptrR* mutant strain has a growth deficiency in the presence of Glu as the sole nitrogen source, suggesting the importance of PtrR-dependent repression of the *yne* operon encoding the GABA-utilization YneI (SSADH) and glutaminase, YneH. It is interesting that a methyl accepting regulator protein (Tar) was upregulated in the *E. coli ptrR* mutant under nitrogen starvation conditions (Table 2).

Extracellular Ptr alters the OmpF porin charge and pore size, resulting in partial pore closure and a consequent decrease in outer membrane permeability (8,35). The transcriptional regulation of *ompF* is known to be regulated by a cascade mechanism involving FnrS and MarA (36,37). PtrR is important for the regulation of the *fnrS* gene as well as the GABA shunt involving the YneI SSADH, the YneH glutaminase, and at least in some enterobacteria, a chemotaxis response-related gene. The *ptrR* null mutant had reduced *ompF and ompL* mRNA during growth in M9 media with glycerol as the carbon source and an L-glutamine supplement, potentially regulated through the FnrS-*marA* regulatory cascade and direct transcriptional regulation of the *yihQPO-ompL* operon. This results in increased resistance to the tetracycline group of antibiotics (i.e., demeclocycline and chlorotetracycline), but not chloramphenicol, erythromycin, and other antibiotics present on Biolog plate 11C (Fig. S2). PtrR thus regulates the *yneIHGF* operon in response to nitrogen and carbon starvation and under Ptr-induced stress conditions in *E. coli*. PtrR transcriptional regulation may explain the Ptr connection with antibiotic resistance development under stress conditions.

## MATERIAL AND METHODS

### Bacterial strains and plasmids

All mutant strains for *yneJ*, *ydiF* and *yneH* were derived from the Keio collection. The parent strain is BW25113 (WT). This reference strain was utilized in growth screens and specified as “wild type”. The strains utilized in the growth screens were first verified using the polymerase chain reaction. The strains for *yneI* promoter LacZ fusions are described in Supplemental Materials.

### Media and growth of cells

All cell cultures were grown overnight in LB media and refreshed the next day for 2 hours. An altered M9 medium was prepared with 10 or 20 mM of Glu and/or Ptr as the nitrogen source and glycerol or glucose as the carbon source. All cultures were diluted to an absorbency at 600 nm of 0.05 in 5 mL of the altered M9 media in culture tubes. Cultures were then transferred for growth detection, or putrescine was added to the growth media to induce putrescine stress. Growth measurement were detected using a BioTek 800 96-well plate reader.

Cultures were grown under the different conditions in culture tubes for an additional 3-5 hours, and 200-400 μl samples were collected and stored frozen at −20℃ for subsequent LacZ assay measurements. The absorbance at 600 nm was measured for each sample in a BIO-RAD SmartSpec Plus Spectrometer.

### RNA sequencing

RNA sequencing data were generated under aerobic exponential growth conditions in M9 medium or under carbon/nitrogen starvation conditions with 0.8% glycerol as the carbon source, with 10 mM putrescine and/or 20 mM glutamate as nitrogen sources or M9 medium supplemented with 10 mM putrescine, 10 mM L-glutamine and 0.8% glycerol as the nitrogen/carbon source. The wild type BW25113 strain was grown as a control for the isogenic *ptrR* mutant strain. Pre-cultures for the RNA sequencing experiments were taken by scraping frozen stocks and growing the cells in LB medium. Cells were washed twice with M9 medium and inoculated at an OD_600_ of 0.05. The cells were collected at an OD_600_ of 0.9 and were harvested using the Qiagen RNA-protect bacterial reagent according to the manufacturer’s specifications. Pelleted cells were stored at −80°C, and after cell resuspension and partial lysis, they were ruptured with a beat beater; the total RNA was extracted using a Qiagen RNA purification kit. After total RNA extraction, the quality was assessed using an Aglient Bioanalyser using an RNA 6000 kit after removal of ribosomal RNA. Paired-end strand specific RNA sequencing libraries were prepared as described (38).

### LacZ enzyme assays

A LacZ buffer with 23 µg/ml of β-mercaptethanol added, referred to as Z-buffer, was prepared for β-galactosidase assays (39,40). All samples were defrosted, and 200 µl of sample was mixed with 800 µl of Z-buffer and 25 µl of chloroform. All tubes were twice vortexed for 10 seconds and then incubated at 37°C for 40-90 min, after which 200 µl of ortho-nitrophenyl-β-galactoside was added. The reaction was stopped with 500 μl of 1M Na_2_CO_3_, and each sample was centrifuged at 13,000 rpm. An aliquot of 200 μl of the supernatant from each sample was transferred to a 96 well microplate, and the absorbance was measured at 405 nm, 450 nm, and 490 nm in a BioTek ELx808 Plate Reader. β-galactosidase activity was measured in Miller units.

### ChIP-exo experiments

The strains harboring 8-myc were generated by a λ red-mediated site-specific recombination system, targeting the C-terminal region as described previously (41). ChIP-exo experimentation was performed following the procedures previously described (42,43), To identify PtrR binding sites for each strain the DNA bound to PtrR from formaldehyde cross-linked cells was isolated by chromatin immunoprecipitation (ChIP) with the antibodies that specifically recognize the myc tag (9E10, Santa Cruz Biotechnology), and Dynabeads Pan Mouse IgG magnetic beads (Invitrogen) were added, followed by stringent washings as described previously. ChIP materials (chromatin-beads) were used to perform on-bead enzymatic reactions of the ChIP-exo method. Briefly, the sheared DNA of the chromatin-beads was repaired by the NEBNext End Repair Module (New England Biolabs), followed by the addition of a single dA overhang and ligation of the first adaptor (5’-phosphorylated) using a dA-Tailing Module (New England Biolabs) and NEBNext Quick Ligation Module (New England Biolabs), respectively. Nick repair was performed by using the PreCR Repair Mix (New England Biolabs). Lambda exonuclease- and RecJf exonuclease-treated chromatin was eluted from the beads, and overnight incubation at 65 degree reversed the protein-DNA cross-link. RNA- and protein-free DNA samples were used to perform primer extension and second adaptor ligation with the following modifications. The DNA samples, incubated for primer extension as described previously (38), were treated with the dA-Tailing Module and NEBNext Quick Ligation Module (New England Biolabs) for second adaptor ligation. The DNA sample, purified using the GeneRead Size Selection Kit (Qiagen), was enriched by polymerase chain reaction (PCR) using Phusion High-Fidelity DNA Polymerase (New England Biolabs). The amplified DNA samples were purified again with a GeneRead Size Selection Kit (Qiagen) and quantified using Qubit dsDNA HS Assay Kit (Life Technologies). The quality of the DNA sample was checked by running the Agilent High Sensitivity DNA Kit using an Agilent 2100 Bioanalyzer before sequencing using HiSeq 2500 (Illumina) following the manufacturer’s instructions. Each modified step was also performed following the manufacturer’s instructions. ChIP-exo experiments were performed in duplicate. For the prediction of the PtrR consensus binding sequence for the *yneI, fnrS* and *yneJ, fnrS* upstream areas, they were downloaded using the PubSEED database for *E. coli* and closely related species. We constructed multiple alignments of the closely related species upstream regions for the genes that correspond to ChIP-exo protected areas. The length of the binding motif was limited to 15 bp using the MEME tool.

**Fluorescent polarization assay (Supplemental Material)**

Table S1. The upregulated genes detected by the RNAseq data for *ptrR* mutant normalized to WT *E. coli* BW25113 strain during growth in M9 medium with L-Glutamate as single nitrogen source and glycerol as the carbon source

Table S2 Biolog plate PM2A area under the growth curve for the growth *ptrR* (*yneJ) E. coli* mutant and WT *E. coli* BW25113 in M9 medium with no carbon source with L-Glutamate added as supplement.

Table S3. Differentially expressed genes for RNAseq data for *ptrR* mutant and WT *E. coli* BW25113 strain during growth in M9 medium with L-glutamine and putrescine as supplement and glycerol as the carbon source.

Figure S1. The *gltS* promoter region for PtrR binding, identified by ChIP-exo.

Figure S2. The Biolog Plate PM2A (at time point 80) for the growth in M9 medium with no primary carbon source but with L-Glutamate added as a supplement **A.** *ptrR E. coli* mutant versus the BW25113 strains **B.** WT *E. coli* BW25113

Fig. S3 The *yihQOP-ompL* operon in Enterobacteria. The gens are marked as arrows numbered: 1-*ompL*, 2-*yihOP* and 3-*yihQ*

Fig. S4 The *marRA* (1, 4) and *marB* (5) chromosomal gene cluster in Enterobacteria. The clusters conserved conserved and include PtrR regulated *ptrR* (8) conserved *yne* (9, 3, 7) operon and *uxaB*-*yneE* (2,15) genes. The additional conserved gene in the cluster is *ydeE* encoding MFS family transporter.

Fig. S5 Fluorescent polarization assay for PtrR binding to the predicted DNA-binding sequence and PhrR as control

## ACKNOWLEDGEMENT

We would like to thank Marc Abrams (UCSD) for editing the manuscript and Dmitry Rodionov for prediction of YneJ regulatory binding sites.

